# G-quadruplex targeting by CX-5461 as a novel therapeutic strategy in testicular germ cell tumors

**DOI:** 10.64898/2026.09.04.749562

**Authors:** Rashidul Islam, Annalena Liesen, C. Saúl Rodríguez Roque, Katrin Paeschke, Hubert Schorle

## Abstract

Testicular germ cell tumors (TGCTs) are highly curable malignancies; however, resistance to cisplatin-based therapy remains a critical clinical challenge. G-quadruplexes (G4s) are specific guanine rich, four-stranded DNA structures that are enriched at distinct genomic regions. Due to their stability, G4s can influence genome stability, gene expression, DNA replication as well as telomere maintenance. G4 stabilization by small molecules has been shown to have anticancer effects in several tumors but remains underexplored in TGCTs. Here, we systematically evaluated the effect of six different G4- ligands (CX-5461, CX-3543, BRACO19, PDS, CM03 and RHPS4) on TGCT cell lines and non-malignant fibroblasts. Among those G4 ligands, CX-5461 showed strongest cellular changes in TGCTs. We observed dose- and time-dependent G4 stabilization upon CX-5461 treatment in TGCT cells, accompanied by impaired RNA polymerase I-dependent transcription and induction of DNA double-strand breaks, whereas only minimal effects were observed in control fibroblasts. These effects led to the activation of p53 signaling, and consequently an upregulation of canonical p53 target genes, including CDKN1A, MDM2, PIDD1, GADD45A, ZMAT3, SESN1 and PPM1D, resulting in G2/M cell-cycle arrest and apoptosis. The presence of potential quadruplex forming sequences (PQSs) in the promoters or gene bodies of these genes suggests that G4 structures may contribute to the p53 transcriptional response. Collectively, these data demonstrate that CX-5461 appears as highly potent G4-ligand inducing p53-dependent molecular cascades leading to cell death in germ cell tumors. Unlike in many other tumors, p53 remains functionally intact in TGCTs, suggesting that CX-5461-mediated activation of the p53 pathway could be selectively targeted as a therapeutic strategy in TGCTs.

## Introduction

Testicular cancer is the most common solid malignancy in men aged 15-44 years, accounting for ∼1-2% of all male tumors (1–3). Its incidence is rising worldwide, with the highest rates in Europe, North America and Oceania (4). Around 95% of cases are Type II testicular germ cell tumors (TGCTs), which arise from developmentally arrested primordial germ cells and are classified into seminomas and non-seminomas (1–3). Seminoma (SE) is the predominant, homogeneous TGCT subtype (∼60%), whereas non-seminomas comprise embryonal carcinoma (EC), choriocarcinoma (CC), yolk-sac tumor (YST), and teratoma (1–3). TGCTs are among the most curable malignancies, with 5-year survival rates of 85-99% depending on the stage (2, 3). The standard care for TGCT consists of radical orchiectomy followed by surveillance, retroperitoneal lymph node dissection, radiotherapy or chemotherapy, depending on histology and stage (2, 3). Despite high cure rates, 3-5% of all TGCTs and 10-15% of metastatic cases develop cisplatin resistance, resulting in relapsed or refractory disease with limited options and poor prognosis (5, 6). Conventional therapies cause substantial acute and long-term toxicities, underscoring the need for alternative treatments (3, 6).

Therapeutic targeting of G-quadruplexes (G4s), four-stranded DNA structures in specific guanine-rich genomic regions. G4s are enriched in regulatory regions and at sites of active DNA replication, where their formation can impede polymerase progression and destabilize the genome (7–18). Because G4 are enriched in at specific locations in the genome including promoters of oncogenes, G4 stabilization by G4 specific small molecules (G4 ligands) can repress the expression of oncogenes (such as *MYC, KRAS* and *BCL2*), modulate *hTERT* and telomerase activity, and induce replication stress and DNA damage, leading to cell-cycle arrest and apoptosis (7–12). Several G4 ligands, including PDS, BRACO-19, RHPS4, CM03, TMPyP4 and Phen-DC3, have demonstrated antitumor activity in preclinical models of various cancers, including breast (13, 19–21), colorectal (22–24), prostate (25, 26), osteosarcoma (27–29), ovarian (14, 15, 30–32), neuroblastoma (33–36), lymphoma (37, 38), glioblastoma (39–41), leukemia (37, 42–46), melanoma (47), pancreatic (48, 49), cervical (12, 50). While CX-3543 (NCT00955786) (51) and APTO-253 (NCT02267863) (52) did not progress beyond early-phase clinical trials, QN-302 (NCT06086522) and CX-5461 (NCT04890613) are in clinical trials for advanced solid tumors and hematological malignancies (53, 54).

The anticancer effects of G4-ligands are attributed to multiple changes such as stabilization of telomeric G4 structures inhibition of RNA polymerase I-dependent ribosome biogenesis, modulation of oncogene expression, induction of genomic instability and synthetic lethality, and interacting with mitochondrial G4s (7–12, 14–18, 54).

To investigate the impact of G4 targeting in TGCTs, here we tested the effects of six G4 ligands (CX-5461, CX-3543, BRACO-19, PDS, CM03 and RHPS4) on cell viability and fate in TGCT cell lines. CX-5461 emerged as the most promising ligand, showing the lowest IC_50_ in TGCT cells and much higher IC_50_ values in control fibroblasts, and it induced G4 stabilization, inhibition of RNA polymerase I transcription, DNA damage, p53 activation, G2/M arrest and apoptosis in TGCT cells, with minimal effects in non-malignant fibroblasts. Genes commonly altered across TGCTs and involved in p53 activation harbored promoter or gene body potential quadruplex forming sequences (PQSs), suggesting that CX-5461 may modulate TGCT cell fate via regulatory G4s and supporting CX-5461 as a promising G4-targeting candidate for TGCT therapy.

## Materials and methods

### Reagents and materials

G4-ligands were obtained from the suppliers listed (supplementary Table S1). Compounds were dissolved in DMSO, except for PDS, which was dissolved in H₂O. Stock solutions (concentrations of 1-10 mM) were stored at -80 °C. Additional reagents used in this study are listed in Supplementary Table S1.

### Cell lines and culture conditions

Human TGCT cell lines TCam-2 (SE), NT2/D1, 2102EP, and NCCIT (EC), JAR and JEG-3 (CC), and GCT72 (YST), were used (55–58). Adult fibroblasts (MPAF) (56–58) and healthy testis fibroblast (Hs1.Tes) served as controls (59). Cells were cultured at 37 °C in a humidified atmosphere containing 7.5% CO₂. Details see supplementary Table S2.

### Cell viability assay

Cells were seeded at 3x10^3^ cells per well in 96-well plates and treated after 24 h with the G4 ligands for 24, 48 or 72 h. For the 72 h condition, compounds were replenished after 48 h. Five replicates were included for each concentration and time point. Cell viability was determined using the XTT assay as described (57–60). IC₅₀ values were determined by nonlinear regression, using an inhibitor vs. response (variable-slope, four-parameter) model in GraphPad Prism.

### Cell cycle analysis

Cell-cycle analysis was performed using propidium iodide (PI) staining, with minor modifications to a previously described protocol (60). Briefly, cells were seeded at 1.5x10^5^ cells per well in 6-well plates, treated with CX-5461 (1 or 2 μM) for 24 h, fixed in 70% ethanol, stained with PI /RNase A, and analyzed on a BD FACSCanto II (Flow Cytometry Core Facility, University Hospital Bonn). Data were analyzed in FlowJo v10.10.

### Apoptosis assay

Apoptosis was assessed using the FITC Annexin V Apoptosis Detection Kit with 7-AAD according to the manufacturer’s protocol. Cells were seeded at 1.5×10^5^ cells per well in 6-well plates and treated with CX-5461 (1 or 2 μM) for 24 or 48 h, stained with FITC Annexin V and 7-AAD, and analyzed on a BD FACSCanto II and data were analyzed in FlowJo.

### Flow cytometric analysis of γ-H2AX

Phosphorylated H2AX (γ-H2AX) was quantified by flow cytometry with minor modifications (61). Cells were seeded at 5×10^4^ cells / mL and treated with CX-5461 (1 or 2 μM) for 24 h or 48 h, stained with Zombie NIR fixable viability dye (BioLegend, # 423105), fixed in 4% PFA, permeabilized in 90% methanol and blocked in 1% BSA. Cells were incubated with anti-γ-H2AX (Ser139) (1:250, 1 h, RT), followed by Alexa Fluor 488-conjugated secondary antibody (1:400, 1 h, RT), then acquired on a BD FACSCanto™ II flow cytometer and data were analyzed using FlowJo. Debris, doublets, and dead cells were excluded by FSC/SSC, FSC-A/FSC-H, and viability gating, respectively (Supplementary Figure 1a). γ-H2AX levels were quantified as the median Fluorescence Intensity (MFI) in live cells, with background levels defined by unstained and secondary-only controls (Supplementary Figure 1b). Fold changes were calculated by normalizing sample MFI to DMSO vehicle controls matched for treatment condition.

### Quantification of G-quadruplex using BG-flow

G4s structures were quantified by flow cytometry using the G4-specific single-chain antibody BG4 (62). Cells (5 ×10^4^ cells / mL) were treated with CX-5461 (1 or 2 μM) for 24 h or 48 h, stained with Zombie NIR fixable viability dye, fixed in methanol/acetic acid (3:1), permeabilized with 0.5% Triton X-100 and incubated with BG4 antibody (1 µg, 2 h) in 10% goat serum followed by anti-FLAG antibody (1:800) in FACS/permeabilization buffer (2:1) containing 10% goat serum for 1h at RT, washed and incubated with Alexa Fluor 488 secondary antibody (1:600; 1h at RT). Samples were acquired on a BD FACSCanto II and analyzed using FlowJo v10.10.0; debris, doublets, and dead cells were excluded by standard gating (Supplementary Figure 2a). BG4 signal (MFI) was quantified in live cells, with background levels defined by unstained and BG4-ve controls (Supplementary Figure 2b). Fold changes were calculated by normalizing sample MFI to DMSO vehicle controls matched for treatment condition.

### 3′ mRNA sequencing and analysis

TCam-2, NT2/D1, 2102EP, JEG3, and GCT72 cells were seeded at 5×10^4^ cells / mL and treated with 2 μM CX-5461 for 24 h. Total RNA was extracted using the Quick-RNA Miniprep Plus Kit. RNA integrity was verified using the RNA ScreenTape system (Agilent). Libraries were prepared with the Watchmaker mRNA Library Prep Kit and sequenced on a NovaSeq 600 platform at the Next Generation Sequencing (NGS) Core Facility (University Hospital Bonn) to yield ∼15 million reads per sample. FASTQ files were processed with nf-core/rnaseq (v3.14.0) using default settings (63). Adapter trimming was performed with Trim Galore, and reads were aligned to the Homo sapiens GRCh38 genome using STAR (v2.7.9a) (64). Gene expression was quantified with Salmon (v1.10.1) (65) and standard quality metrics (FastQC, alignment statistics, gene body coverage) were generated automatically. Sorted and indexed BAM files were processed in R (v4.5.2) with Rsamtools, and gene counts were obtained using featureCounts (Rsubread v2.12.0) (66) and Ensembl GRCh38.114 annotation, excluding multi-mapping reads. Differential expression analysis was performed using DESeq2 (v1.38.3) (67) after filtering genes with total counts < 10. Log2 fold changes were shrunk with apeglm (v1.20.0), and genes with log2FC > 0.58 and FDR < 0.05 were considered significant. Functional annotations were derived from MSigDB Hallmark (msigdbr v7.5.1) (68) and KEGG (69). Gene Set Enrichment Analysis (GSEA) was performed using clusterProfiler (v4.6.0) for KEGG and MSigDB Hallmark gene sets (parameters: p < 0.05, q < 0.2, minGSSize = 10, maxGSSize = 500) (70). Results with FDR < 0.05 were considered significant. All statistical tests were two-sided and corrected for multiple comparisons using the Benjamini-Hochberg method. Data visualizations were generated with ggplot2, ggrepel, and ComplexHeatmap, and Venn diagrams were created with the online tool InteractiVenn (71).

### PQSs prediction and scoring using pqsfinder

Promoter- and gene-body G-quadruplex (G4) propensity was assessed in seven p53 target genes, namely CDKN1A, MDM2, PIDD1, GADD45A, ZMAT3, SESN1, and PPM1D, using the human reference genome GRCh38/hg38. Gene coordinates were obtained from TxDb.Hsapiens.UCSC.hg38.knownGene (version 3.22.0), and gene symbols were mapped to Entrez Gene identifiers using org.Hs.eg.db (version 3.22.0) to ensure compatibility with the annotation framework. Promoter regions were defined as 1,000 bp upstream and 200 bp downstream of the transcription start site, whereas gene bodies encompassed the full transcribed sequence from transcription start to termination site. Putative G4 structures were predicted using pqsfinder (version 2.26.0) (72), with a minimum score threshold of 47, a maximum PQSs length of 50 nucleotides, a minimum length of 12 nucleotides, and overlapping hits enabled (72, 73). To capture strand-specific patterns, sequence scans were performed independently on both DNA strands for each genomic interval.

### RT-qPCR

Total RNA was isolated using the Quick-RNA Miniprep Plus Kit and treated with DNase I according to the manufacturer’s instructions. cDNA synthesis and SYBR Green-based RT-qPCR were performed as previously described (56). Primers are listed (Supplementary Table S3). Relative gene expression levels were normalized to the mean Ct of GAPDH and ACTIN and calculated using the ΔΔCt method.

### Protein extraction, SDS-PAGE and western blot

Cells were lysed in RIPA buffer supplemented with protease inhibitors (Roche), sonicated, and centrifuged (13,000 x g) 15 min, 4 °C). Protein concentration was measured using NanoDrop One. Proteins were separated by SDS-PAGE and transferred to PVDF membranes using the Trans-Blot Turbo system, as described previously (56). Membranes were probed with antibodies against phospho-H2AX (Ser139) (1:1000) and α-tubulin (1:1000), followed by HRP-conjugated secondary antibodies (1:2000). Signals were detected using ECL (WESTAR NOVA 2.0) on a ChemiDoc MP system. Chemiluminescent signals were detected with the ChemiDoc MP imaging system (Bio-Rad Laboratories) using WESTAR NOVA 2.0 ECL substrate.

### Figure preparation and statistical analysis

Graphs were generated using GraphPad Prism version 10.6.1 (GraphPad Software) and assembled using Inkscape version 1.4.3. Data represents standard deviation (SD) from independent biological replicates. Statistical significance between two groups was determined using an unpaired, two-tailed Student’s t-test. Significance levels are indicated as follows: ns, not significant; *P < 0.05; **P < 0.001; ***P < 0.0001; ****P < 0.00001.

### Data availability

The 3′ mRNA sequencing data generated from TGCT cell lines in this study will be deposited in the Gene Expression Omnibus (GEO) database, and the accession code will be provided in the final version of the manuscript.

## Results

### CX-5461 markedly reduces the viability of TGCT cells

To assess the impact of G4 ligands on TGCT cell survival, we tested six different, well characterized, G4 ligands (CX-5461, CX-3543, BRACO-19, PDS, CM03 and RHPS4) on their effect in TGCT subtype-specific cell lines and control fibroblasts. To reveal metabolic changes upon treatment and determine IC_50_ values, we performed XTT assays after exposing cells to increasing ligand concentrations for 24, 48 and 72 h to examine concentration- and time-dependent effects and derive IC₅₀ values for each compound (Table-1; Supplementary Table S4-S8).

CX-5461 exhibited significant dose-and time-dependent cytotoxicity in all TGCT cell lines (except GCT72) achieving IC₅₀ values of 0.1-1.6 µM at 72 h (Figure 1a-g; Table-1; Supplementary Figure 3a-g). Across time points, IC₅₀ values were similar among TGCT lines, with NT2/D1 and NCCIT being most sensitive at 72 h, whereas GCT72 had displayed higher IC₅₀ values (Figure 1a-g; Table 1; Supplementary Figure 3a-f). Control fibroblasts MPAF and Hs1.Tes showed minimal response up to 48 h and responded only at 72 h, with the highest IC₅₀ values of all cell types (Figure 1h-i; Table-1; Supplementary Figure 3h-i).

**Figure 1:**
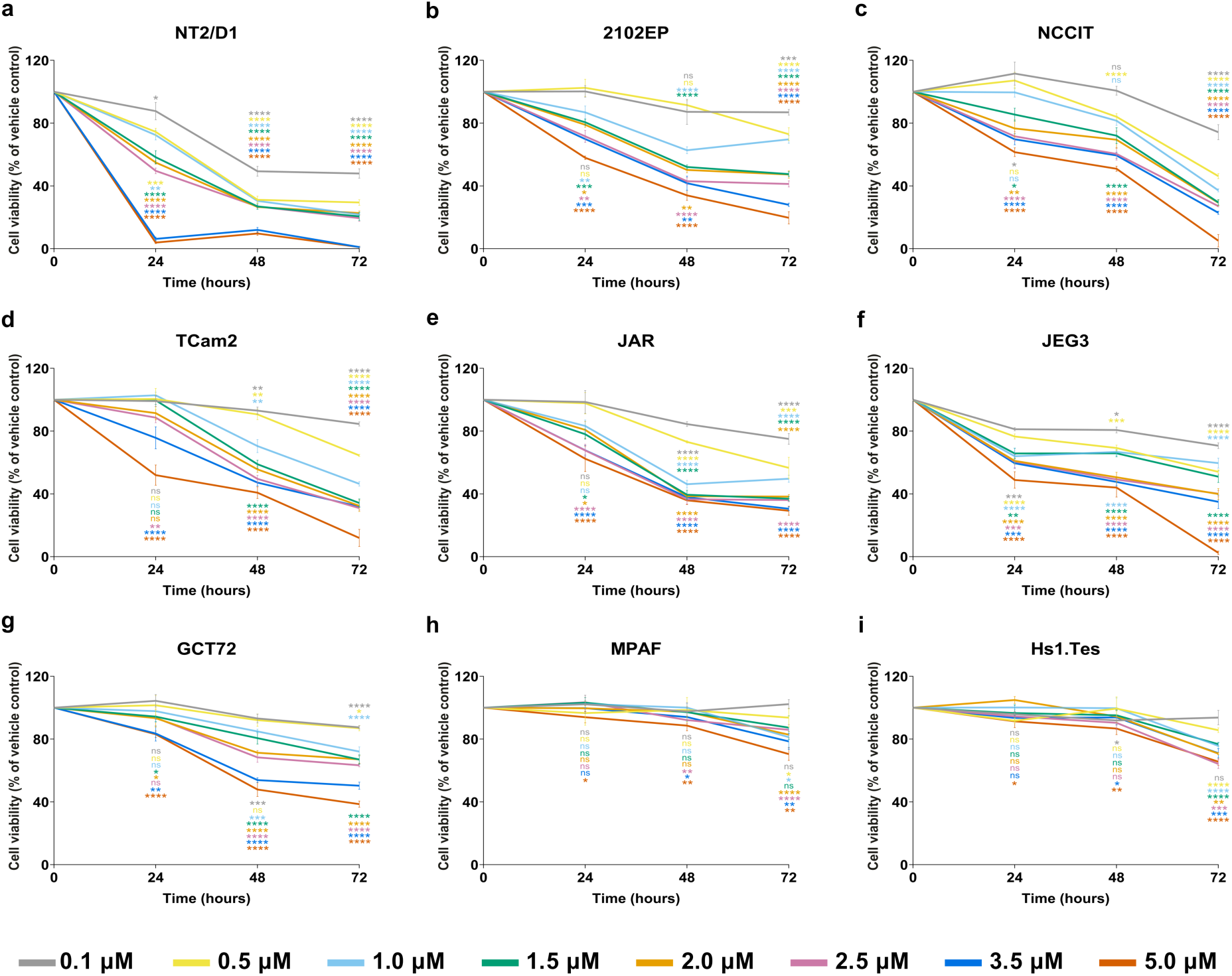
CX-5461 treatment significantly reduces TGCT cell viability in a dose- and time-dependent manner. XTT cell viability assays (n = 5) were performed on TGCT **(a-g)** and non-malignant control cell lines treated with 0.1, 0.5, 1.0, 1.5, 2.0, 2.5, 3.5, and 5.0 µM CX-5461 for 24, 48, and 72 hours. CX-5461 exposure resulted in a significant, concentration- and time-dependent decrease in TGCT cell viability, whereas MPAF and Hs1.Tes control fibroblasts **(h-i)** were comparatively less affected. DMSO vehicle controls were included at each concentration and set to 100% viability; data are normalized to the corresponding vehicle controls and presented as mean ± SD. Statistical significance was assessed using a two-tailed Student’s t-test. Asterisks indicate statistical (*P < 0.05, **P < 0.01, ***P < 0.001, ****P < 0.0001), and non-significant differences are denoted as ns (P > 0.05); asterisk color denotes the corresponding CX-5461 concentration.

**Table 1:** IC50 values (µM) for TGCT and control cell lines following CX-5461 treatment, determined using the XTT viability assay. . Results are presented for 24, 48, and 72 h post-treatment. Color coding denotes potency: red (0-2 µM, very high potency), blue (>2-6 µM, high potency), Yellow (>6-12 µM, moderate potency), and Teal (>12 µM-Unstable, low potency).

| Entity | Cell line | IC50 (μM) |  |  |
| --- | --- | --- | --- | --- |
|  |  | 24 h | 48 h | 72 h |
| Embryonal carcinoma | NT2/D1 | 1.775 | 0.1037 | 0.1003 |
|  | 2102EP | 6.586 | 2.097 | 1.584 |
|  | NCCIT | 6.202 | 5.065 | 0.4554 |
| Seminoma | TCam2 | 5.137 | 2.797 | 0.8702 |
| Choriocarcinoma | JAR | 8.003 | 1.209 | 0.7988 |
|  | JEG3 | 6.739 | 3.055 | 0.9163 |
| Yolk sac tumor | GCT72 | 15.31 | 4.506 | 3.708 |
| Fibroblast | MPAF | Unstable | 26.02 | 20.06 |
|  | Hs1.Tes | Unstable | 42.27 | 16.10 |

BRACO19 exerted dose-dependent cytotoxicity across all TGCT cell lines, with very high potency (IC₅₀ 0-16 µM) at 72 h (Supplementary Figure 4a-g; Supplementary Table S4). At 24 h and 48 h, IC₅₀ values indicated high (IC₅₀ >16-35 µM) to moderate (IC₅₀ >35-72 µM) cytotoxicity in all TGCT cell lines, and MPAF showed a similar dose- and time-dependent profile (Supplementary Figure 4a-h; Supplementary Table S4).

CM03 induced strong dose- and time-dependent cytotoxicity across TGCT cell lines, with very high potency at 72 h (IC₅₀ 0-0.2 µM) in all cell lines except GCT72 and the lowest IC₅₀ (0.0001 µM) in TCam2 (Supplementary Figure 5a-g; Supplementary Table S5). In contrast, MPAF remained largely insensitive up to 48 h and responded only after 72 h, still with higher IC₅₀ values than any TGCT line (Supplementary Figure 5h; Supplementary Table S5).

PDS elicited a heterogeneous cytotoxic response across TGCT cell lines, with high potency at 72 h in JAR, JEG3, NT2/D1 and NCCIT (IC₅₀ 0-10 µM; lowest <3.3 µM in JAR and JEG3) and lower sensitivity in TCam2 and GCT72 (IC₅₀ >10-20 µM) (Supplementary Figure 6a-g; Supplementary Table S6). PDS induced a similar level of cytotoxicity in MPAF (Supplementary Figure 6h; Supplementary Table S6).

RHPS4 induced clear dose- and time-dependent cytotoxicity in TGCT cell lines. After 48 h and 72 h, EC and CC lines were highly sensitive (IC₅₀ 0-3 µM), whereas TCam2 showed high sensitivity (IC₅₀ 3-10 µM) (Supplementary Figure 7a-f; Supplementary Table S7). GCT72 exhibited only moderate effects (IC₅₀ >10-20 µM) up to 48 h, and viability unexpectedly increased at 72 h, precluding reliable IC₅₀ determination and suggesting transient or adaptive responses (Supplementary Figure 7g; Supplementary Table S7). MPAF were minimally affected at 24 h and 48 h but responded at 72 h (IC₅₀ >3-10 µM), indicating limited TGCT selectivity at later time points (Supplementary Figure 7h; Supplementary Table S7).

Lastly, CX-3543 induced very high cytotoxicity (IC₅₀ 0-4 µM) in all TGCT cell lines and in MPAF at 24, 48 and 72 h, with TCam2 being most sensitive (Supplementary Figure 8a-h; Supplementary Table S8). Notably, JEG3 and MPAF appeared to partially regain viability after 72 h of exposure (Supplementary Figure 8f, h; Supplementary Table S8).

Based on the observed sensitivity and response profiles, the G4 ligands were ranked by preferential cytotoxicity toward TGCT cells over control fibroblasts as- CX-5461, CM03, RHPS4, PDS, BRACO19 and CX-3543. Hence, we selected CX-5461 for further detailed analysis.

### CX-5461 induces dose- and time-dependent G4 stabilization in TGCT cells

To investigate G4 specific formation/ accumulation upon CX-5461 treatment, we treated TGCT cell lines and control fibroblasts with 1 or 2 µM CX-5461 for 24 or 48 h, at concentrations within the IC30-IC50 range for TGCT cells, and quantified G4 levels by BG4-flow cytometry (62). CX-5461 induced a robust, dose- and time-dependent increase in BG4 signal in TGCT cell lines, detectable at 24 h and further increased at 48 h, with the extent of G4 induction differing between TGCT subtypes (Figure 2a-f). In NT2/D1, 2102EP and NCCIT cells, BG4 signal was ∼2.2-3.8-fold increased after 2 µM for 48 h, whereas TCam2, JAR and JEG3 showed ∼1.5–2.5-fold increases (Figure 2a-f). In contrast, MPAF showed no detectable increase in BG4 signal at the selected concentrations or time points (Figure 2g).

**Figure 2:**
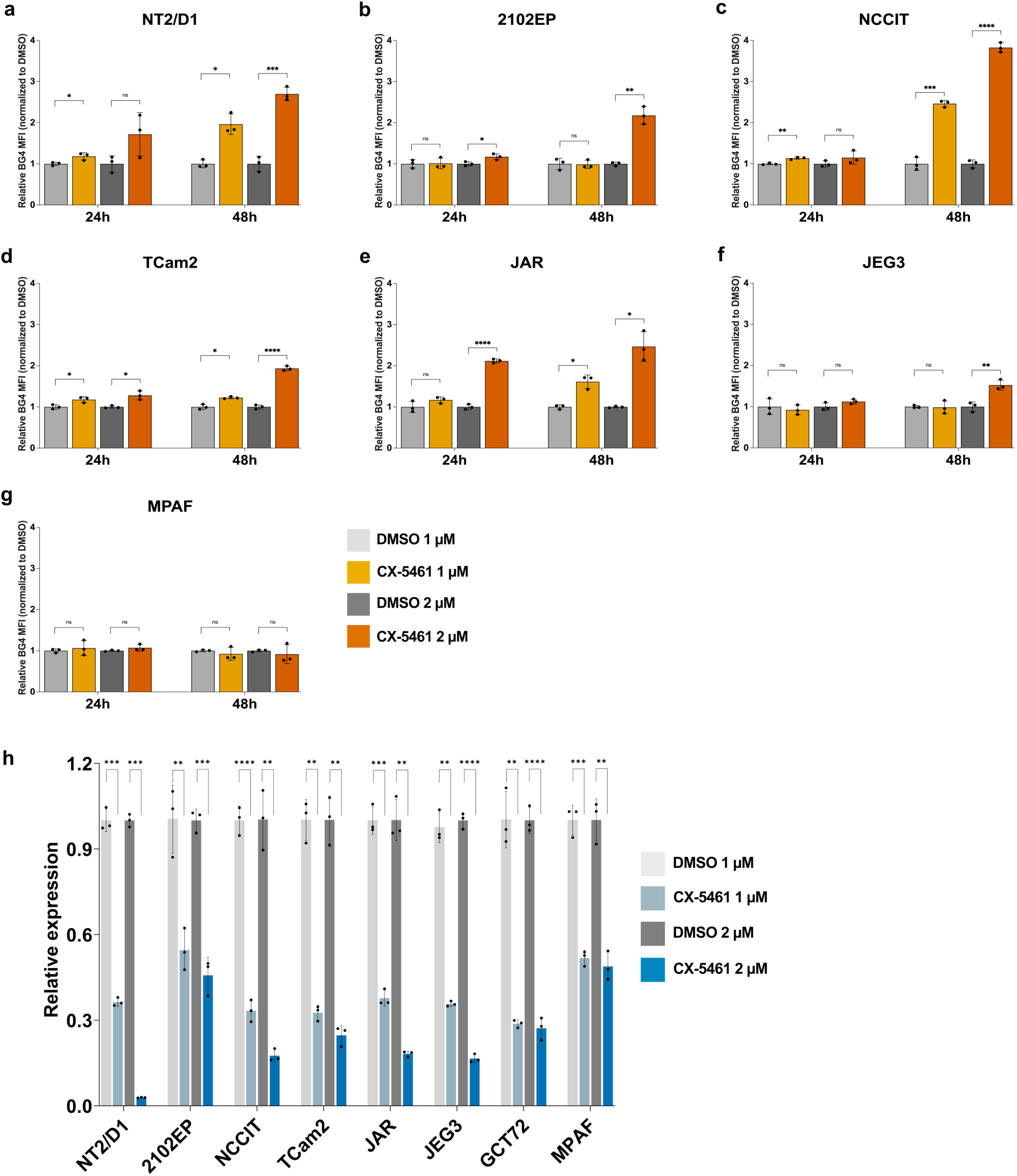
CX-5461 significantly enhances BG4 signal and reduces 47S pre-rRNA expression in TGCT cells. TGCT cell lines **(a-f)** and control MPAF fibroblasts **(g)** were treated with 1 µM or 2 µM CX-5461 for 24 h or 48 h. BG4 signal intensity was quantified relative to corresponding DMSO controls (n = 3 independent experiments), with data presented as mean ± SD. CX-5461 significantly increased BG4 signal in TGCT cells in a dose- and time-dependent manner, whereas minimal or no changes were observed in MPAF fibroblasts. **(h).** Relative 47S pre-rRNA expression levels were measured by RT-qPCR in TGCT cell lines and MPAF cells treated with DMSO or CX-5461 (1 or 2 µM) for 1 h. Expression was normalized to GAPDH and β-actin and presented relative to DMSO controls. Data are shown as mean ± SD from independent experiments (n = 3). Statistical analysis was performed using a two-tailed Student’s t-test relative to DMSO (*P < 0.05, **P < 0.01, ***P < 0.001, ***P < 0.0001; ns, not significant, P > 0.05).

### CX-5461 reduces 47S pre-rRNA expression in TGCT cells

As CX-5461 has been shown to inhibit RNA polymerase I activity in several cancers (10, 12, 15), we next assessed its effect on rRNA transcription in TGCT cells. TGCT cell lines and MPAF cells were treated with 1 or 2 µM CX-5461 to assess the initial inhibition of 47S pre-rRNA expression, by RT-qPCR. and early inhibition of 47S pre-rRNA expression was quantified by RT-qPCR. CX-5461 induced a clear, dose-dependent reduction of 47S pre-rRNA expression in all TGCT cell lines, most pronounced in NT2/D1 (Figure 2h). In contrast, MPAF showed relatively low decrease of 47S pre-rRNA (Figure 2h), indicating that CX-5461 more strongly suppresses 47S pre-rRNA in TGCT cells under these conditions.

### CX-5461 induces DNA double-strand breaks in TGCT cells

Given that CX-5461-induced G4 stabilization can trigger DNA damage (17, 18), we next quantified γH2AX, a marker of the DNA damage, by flow cytometry in TGCT cell lines and control fibroblasts treated with 1 or 2 µM CX-5461 for 24 or 48 h. CX-5461 caused a clear dose- and time-dependent γH2AX increase in TGCT lines (Figure 3a-h). NT2/D1, 2102EP and NCCIT mounted the strongest response, with ∼2.7-4.9-fold higher γH2AX after 48 h at 2 µM, corroborated by western blot in NT2/D1 (Figure 3b-d; Supplementary Figure 2c). GCT72 showed similarly high damage (∼4-fold), whereas TCam2, JAR and JEG3 displayed more modest but still significant γH2AX induction (Figure 3e-h). In contrast, MPAF showed no change in γH2AX under the same conditions (Figure 3i), indicating CX-5461-induced DNA damage in TGCT but not control fibroblasts.

**Figure 3:**
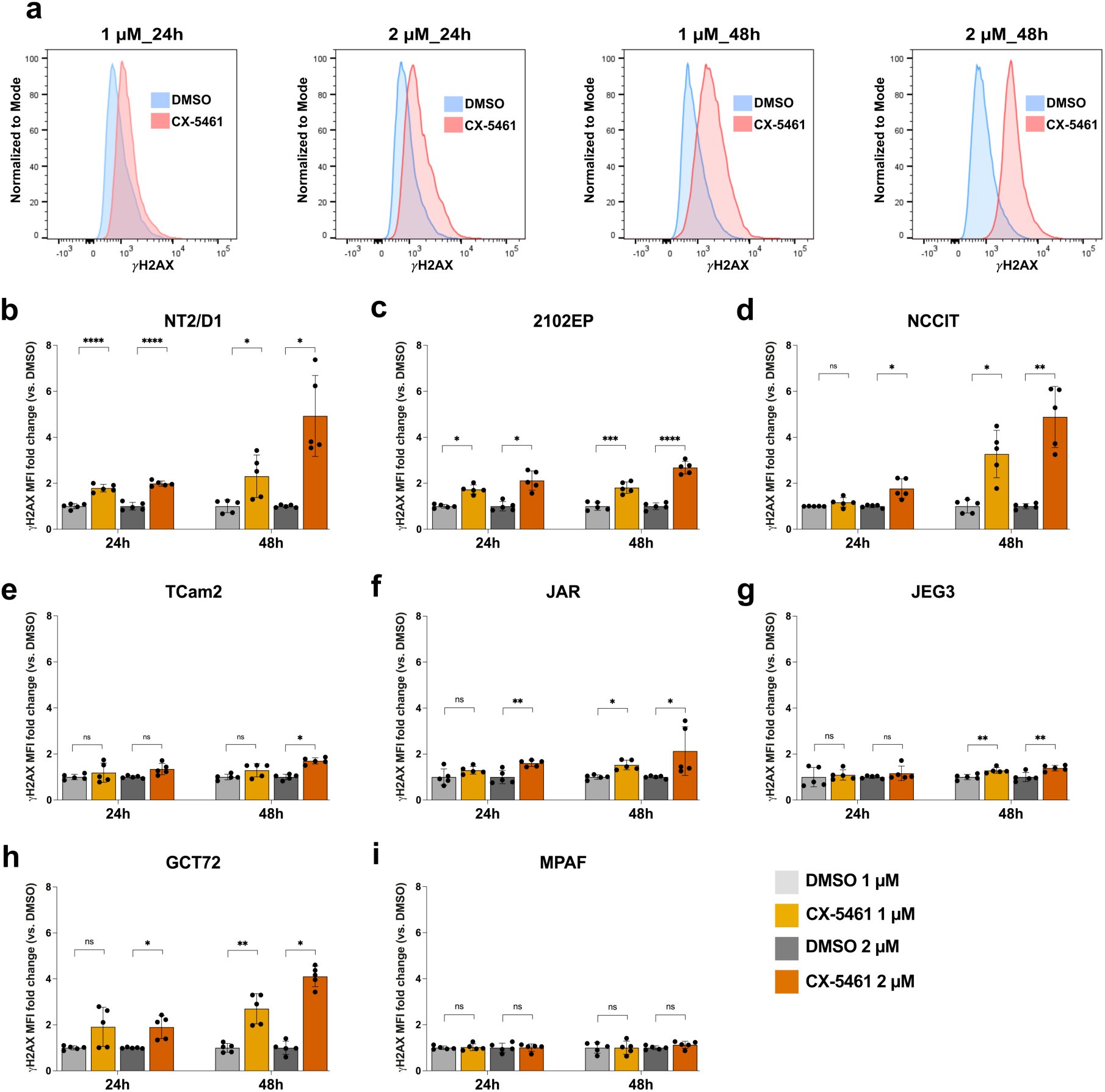
CX-5461 induces double-strand DNA breaks in TGCT cells as detected by γH2AX flow cytometry. TGCT cell lines and non-malignant control cells were treated with CX-5461 (1 and 2 µM) for 24 h and 48 h, followed by flow cytometric analysis of γH2AX levels (n = 5 independent experiments; data are shown as mean ± SD). **a.** Representative γH2AX histograms of NT2/D1 cells treated with CX-5461 at the indicated concentrations and time points compared with corresponding DMSO controls. Quantification of CX-5461-induced double-strand DNA breaks, as reflected by increased γH2AX signal. CX-5461 treatment significantly increased DNA damage in TGCT cells **(b-h)** in a dose- and time-dependent manner, whereas control cells MPAF **(i)** showed no or only minimal changes. Statistical analysis was performed using a two-tailed Student’s t-test relative to DMSO (*P < 0.05, **P < 0.01, ***P < 0.001, ****P < 0.0001; ns, not significant (P > 0.05).

### CX-5461 induces G2/M cell-cycle arrest in TGCT cell lines

Following the observed induction of γH2AX, indicative of a DNA-damage response that can activate cell-cycle checkpoints, we next assessed the effects of CX-5461 on cell-cycle progression in TGCT cells. Cell-cycle analysis after 24 h of treatment with 1 or 2 µM CX-5461 showed a dose-dependent accumulation of TGCT cells in G2/M, increasing from approximately 20-33% in DMSO-treated cultures to 37-77% at 1 µM and 45-85% at 2 µM, with corresponding reductions in G1 and S-phase fractions across all TGCT lines, with the strongest arrest in NCCIT, NT2/D1 and GCT72 cells, followed by JEG3 and TCam2. By contrast, MPAF showed minimal changes in cell-cycle distribution under the same conditions (Figure 4), indicating that CX-5461 elicits a dose-dependent G2/M arrest across TGCT subtypes while sparing non-malignant control cells.

**Figure 4:**
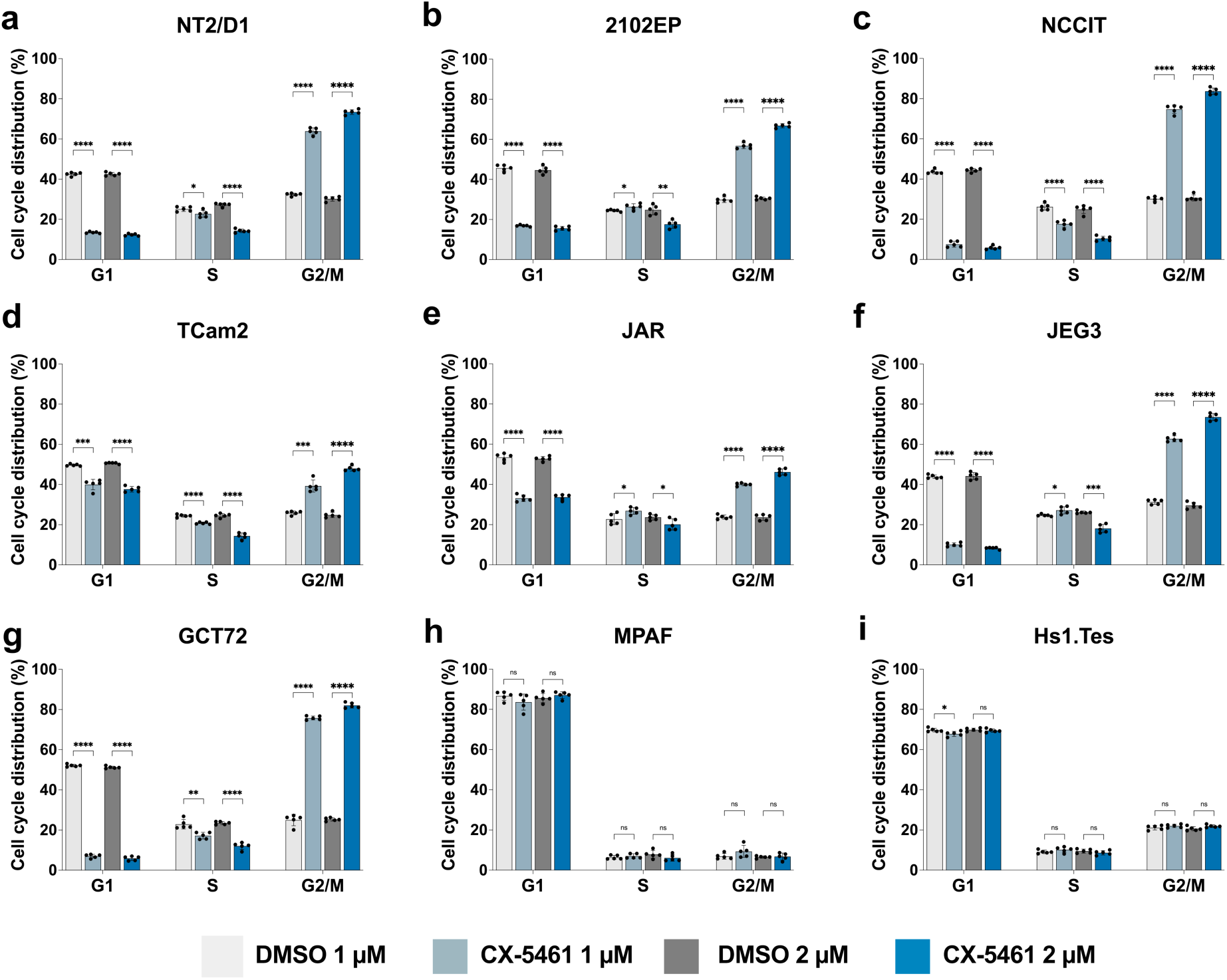
CX-5461 induces G2/M cell cycle arrest in TGCT cells. TGCT cell lines **(a-g)** and non-malignant MPAF and Hs1.Tes fibroblasts **(h-i)** were treated with CX-5461 (1 and 2 µM) for 24 h, followed by propidium iodide (PI) staining and flow cytometric analysis of cell cycle distribution (n = 5 independent experiments; data are shown as mean ± SD). CX-5461 treatment induced a significant, dose-dependent accumulation of TGCT cells in the G2/M phase, whereas cell cycle profiles of control fibroblasts remained largely unchanged. Statistical comparisons for each phase were performed against DMSO-treated controls using a two-tailed Student’s *t*-test. Asterisks indicate statistical significance (*P* < 0.05, \**P* < 0.01, \*\**P* < 0.001, \*\*\**P* < 0.0001); *ns* denotes non-significance (*P* > 0.05).

### CX-5461 triggers apoptosis in TGCT cells in a dose- and time-dependent manner

To determine the downstream consequences of CX-5461-induced DNA damage and G2/M accumulation, we next assessed whether CX-5461 treatment induced apoptosis. TGCT cells and control fibroblasts were treated with 1 or 2 µM CX-5461 for 24 or 48 h, and apoptosis was quantified by Annexin V/7-AAD flow cytometry (Figure 5a). In NT2/D1, 2102EP and NCCIT, total apoptosis increased from ∼10% in DMSO to ∼45-55% after 48 h at 2 µM (Figure 5b-d). TCam2 and JEG3 reached ∼30-36% total apoptosis, with late apoptosis dominating in JEG3 and a higher proportion of early apoptosis in TCam2 at 2 µM, 48 h (Figure 5e, f). GCT72 mounted the weakest response but still showed a clear dose- and time-dependent increase in apoptosis (Figure 5g). In contrast, total apoptosis in MPAF and Hs1.Tes cells remained below 10% under the same conditions and was comparable between DMSO and CX-5461 treatments (Figure 5h-i).

**Figure 5:**
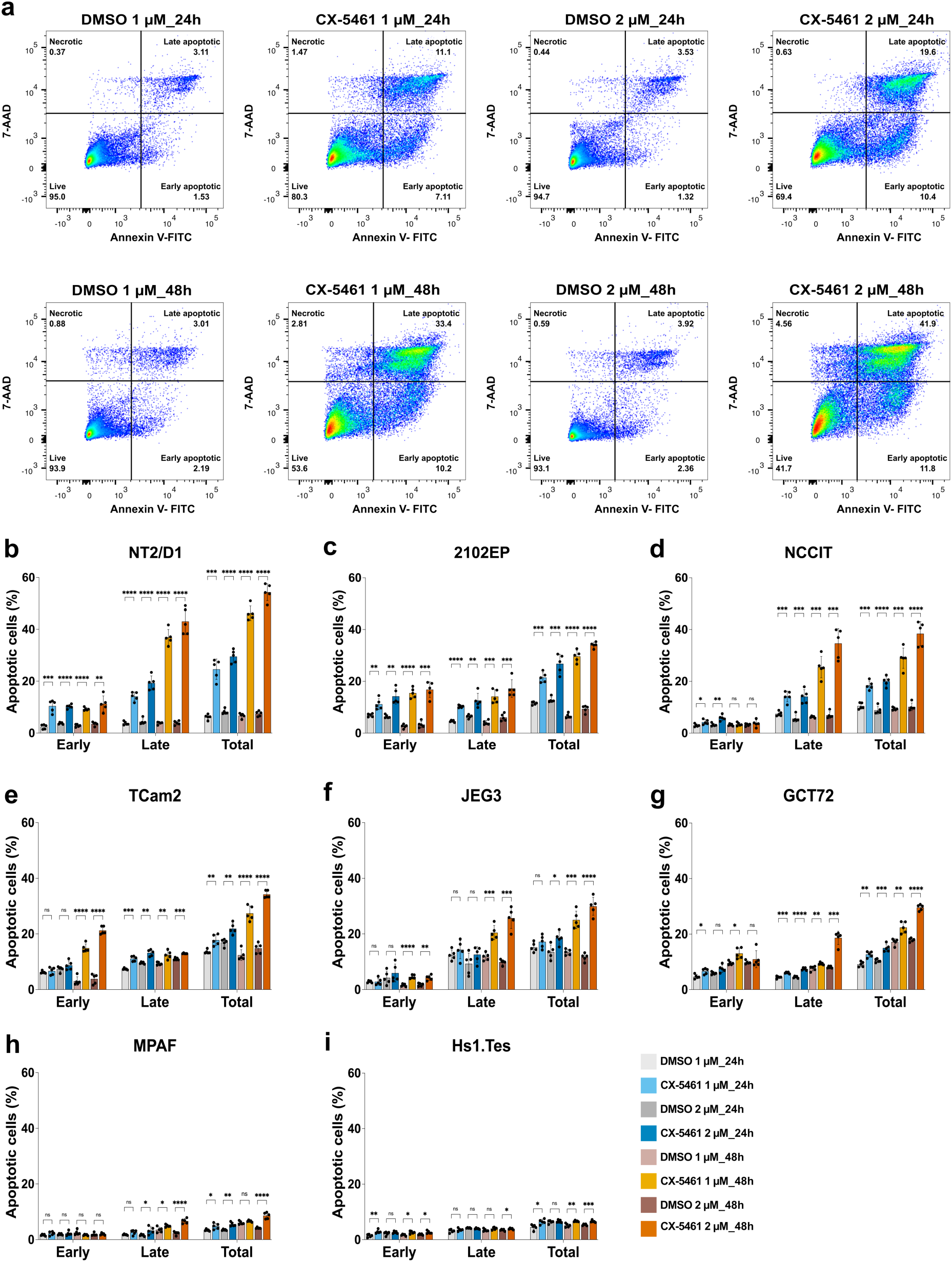
CX-5461 induces dose- and time-dependent apoptosis in TGCT cells. Flow cytometry-based apoptosis analysis using FITC-Annexin V and 7-AAD staining was performed in TGCT cell lines, MPAF and Hs1.Tes fibroblasts cells treated with CX-5461 (1 or 2 µM) for 24 h or 48 h. **a**. Representative flow cytometric plots showing that CX-5461 treatment markedly increased the proportion of apoptotic NT2/D1 cells, with distinct distributions of early and late apoptosis across doses and time points. Quantification of early and late apoptotic populations in TGCT cell lines **(b-g)** as well as in non-malignant MPAF and Hs1.Tes cells **(h-i)** (n = 5 independent experiments; data are shown as mean ± SD). CX-5461 treatment significantly increased apoptotic cell fractions in TGCT lines in a dose- and time-dependent manner, whereas only minimal apoptosis was observed in control fibroblasts. Statistical analysis was performed using a two-tailed Student’s t-test relative to DMSO (*P < 0.05, **P < 0.01, ***P < 0.001, ****P < 0.0001; ns, not significant (P > 0.05).

### CX-5461 activates the p53 pathway in TGCT cell lines, and shared p53 pathway genes harbor promoter and gene body G4s

To examine the transcriptional changes of CX-5461 treatment, we performed bulk RNA-seq in TCam2, NT2/D1, 2102EP, JEG3 and GCT72 cells treated with 2 µM CX-5461 or DMSO for 24 h. Differential expression analysis identified 1,033 significantly differentially expressed genes (DEGs) in TCam2 (711 upregulated, 322 downregulated), 2,934 in NT2/D1 (2,033 upregulated, 901 downregulated), 787 in 2102EP (693 upregulated, 94 downregulated), 3,901 in JEG3 (2,100 upregulated, 1,801 downregulated) and 541 in GCT72 (459 upregulated, 82 downregulated), using thresholds of log2FC > 0.58 and FDR < 0.05 (Supplementary Table S9-S13). Pathway enrichment analysis using KEGG pathways and MSigDB Hallmark gene sets (parameters: p < 0.05, q < 0.2, minGSSize = 10, maxGSSize = 500) were performed, and the top 20 pathways across TGCT cell lines were visualized (Figure 6a,c; Supplementary Table S14-S23). Across TGCT cell lines, p53 signaling was the dominant activated pathway in both KEGG and Hallmark analyses (Figure 6a, c). Within this pathway, 12 genes in the KEGG analysis and 21 genes in the p53-related Hallmark set were enriched across all TGCT cell lines, and intersection analysis identified seven common upregulated p53-responsive genes (CDKN1A, MDM2, PIDD1, GADD45A, ZMAT3, SESN1 and PPM1D) following CX-5461 treatment (Figure 6b-f). These findings were further validated by qPCR analysis in TGCT cell lines (Figure 6g-i).

**Figure 6:**
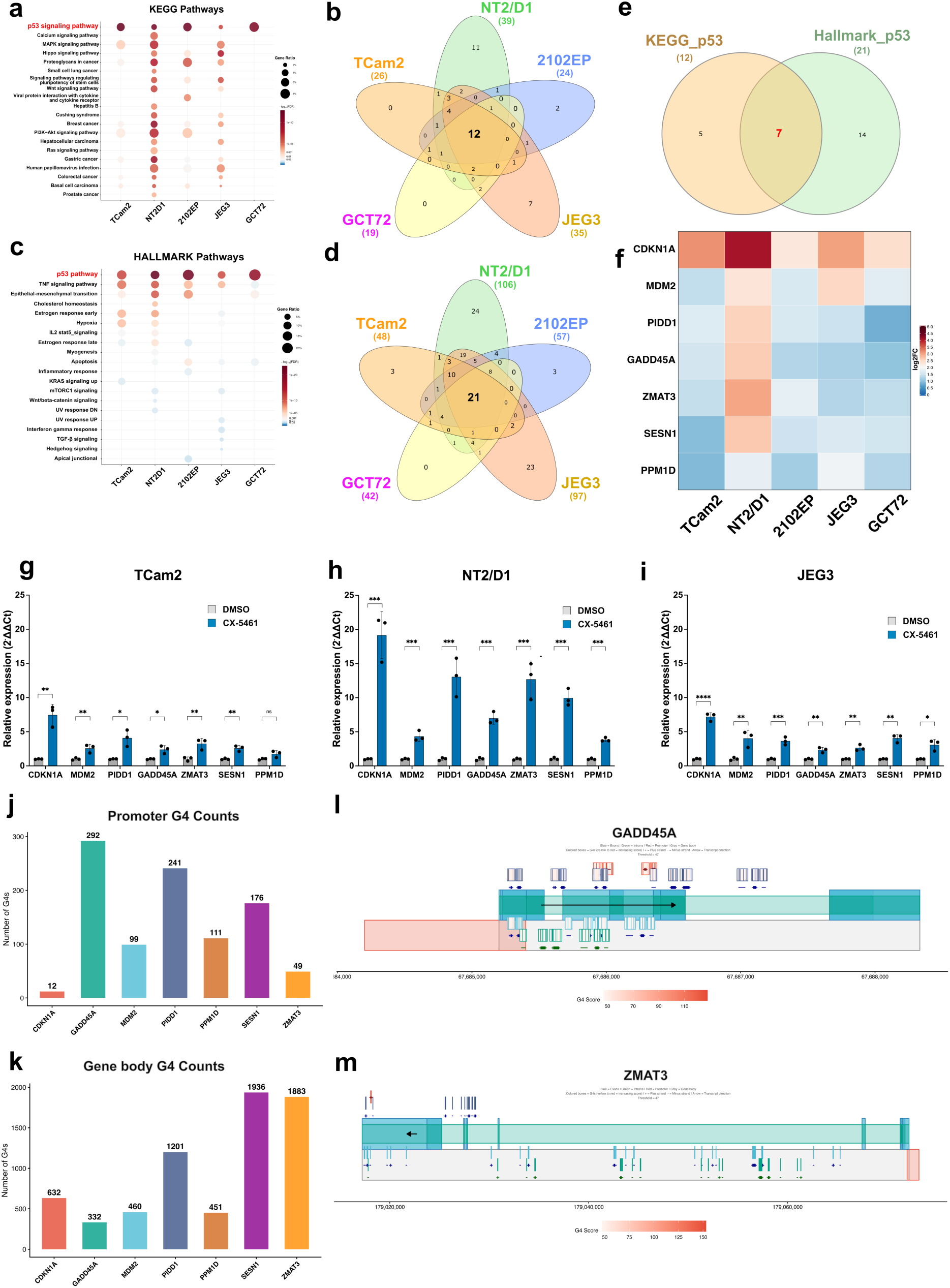
CX-5461 activates p53 pathways, and predicted G4s landscapes in p53 target genes. **a**. Top 20 enriched signaling pathways identified by KEGG analysis across TGCT cell lines**. c.** Top 20 enriched signaling pathways identified by Hallmark gene set analysis across TGCT cell lines. **b.** Venn diagram showing common and unique genes enriched in the p53 signaling pathway (KEGG) in each TGCT cell line. **d.** Venn diagram showing common and unique genes enriched in the p53 signaling pathway (MSigDB Hallmark) in each TGCT cell line. **e.** Overlap of genes enriched in the p53 pathway in both KEGG and MSigDB Hallmark analyses across all TGCT cell lines. **f.** Heatmap displaying log2 fold changes from 0 to +5 using a blue-white-red color gradient for p53 pathway genes common to KEGG and MSigDB Hallmark analyses**. g-i.** RT-qPCR validation of common gene expression (n = 3, data are present as mean ± SD) in TGCT cells treated with CX-5461 compared to DMSO, normalized to the average CT value GAPDH and β-actin. Statistical analysis was performed using a two-tailed Student’s t-test relative to DMSO (*P < 0.05, **P < 0.01, ***P < 0.001, ****P < 0.0001; ns, not significant (P > 0.05). Predicted G4 architecture in CX-5461-responsive p53 target genes. Number of pqsfinder-predicted putative G4-forming sequences (PQSs) in promoter (**j**) and gene body (**k**) regions of seven common p53 target genes (PQSs above the default pqsfinder minimum score threshold of 47). Schematic maps of promoter G4s in GADD45A (**l**) and gene-body G4s in ZMAT3 (**m**).

Next, we used pqsfinder (72, 73) to predict putative G4-forming sequences in the promoter (Figure 6j) and gene-body (Figure 6k) regions of p53 target genes that were consistently upregulated following CX-5461 treatment across TGCT cell lines. Promoter and gene-body regions were selected because G4 structures at these loci can influence transcription and replication-fork progression, enabling the identification of candidate G4-forming loci in CX-5461-responsive p53 target genes. We quantified the G4s per gene, cataloguing PQSs by genomic position, strand, length, sequence, guanine content, and pqsfinder score (Supplementary Table S24). All seven genes harbored predicted G4s in both their promoter and gene body. Across promoters, we identified 12-292 PQSs per gene above the default pqsfinder minimum score threshold of 47 (Figure 6j), with GADD45A showing the highest promoter G4 burden (292 PQSs; Figure 6j). Promoter pqsfinder scores ranged from 74 to 128 across the genes (Supplementary Table S24), with the top-scoring promoter G4s in GADD45A, indicating that promoters of the target genes harbor high-confidence G4 candidates.

Gene bodies were even more densely populated with predicted G4s, with 332-1936 PQSs per gene above the same score threshold (Figure 6k) and SESN1 exhibiting the highest gene-body burden (1936 PQSs) (Figure 6k), consistent with its larger gene body span (∼116 kb). However, G4 density did not simply track gene length: ZMAT3 (∼19 kb) had the second-highest number of gene body PQSs (1883), whereas PPM1D (∼64 kb) contained only 451 gene body PQSs, underscoring that G4 burden is driven by local guanine-rich sequence content rather than gene size alone. The highest pqsfinder scores in gene body PQSs across these genes ranged from 89 to 153 (Supplementary Table S24), with the top-scoring gene body G4s observed in ZMAT3, suggesting that these p53 target genes contain highly G4-prone regions within their transcribed sequences that could influence transcriptional elongation and RNA processing. Comparison of promoter and gene-body regions revealed that GADD45A harbored its highest-scoring predicted PQS in the promoter, at chr1:67,685,983-67,686,014 on the negative strand (32 bp). By contrast, the highest-scoring PQSs of the remaining six genes were located within gene bodies. Among these, ZMAT3 exhibited the highest-scoring gene body PQS, located at chr3:179,057,477-179,057,499 on the positive strand (23 bp) (Supplementary Table S24). Schematic positions of promoter G4s in GADD45A and gene-body G4s in ZMAT3 are shown in Figure 6l-m.

Together, these data show that all seven CX-5461-responsive p53 target genes harbor PQSs in both promoter and gene body regions, suggesting that these G4s may influence CX-5461-dependent transcriptional responses.

## Discussion

In this study, we evaluated six G4 ligands (CX-5461, PDS, CX-3543, CM03, RHPS4 and BRACO-19) in TGCT cell lines. Based on the sensitivity and response profiles for TGCT cells, CX-5461 was further investigated. CX-5461 induces dose- and time-dependent G4 stabilization, RNA polymerase I (Pol I) inhibition and DNA double-strand damage, leading to activation of the p53 signaling pathway. This cascade resulted in G2/M phase arrest and ultimately induced apoptosis across TGCT cell lines (most prominently in EC cells), while non-malignant fibroblasts were only mildly affected. Notably, the common p53 pathway genes across all cell lines harbored strong, stable PQSs in both their promoter and gene body regions, suggesting a potential role for G4-mediated regulation in shaping CX-5461-responsive transcriptional programs in TGCTs.

We found that all six ligands display distinct sensitivity and response profiles across TGCT cell lines and fibroblasts, largely mirroring their reported activities in other cancers. After 72 h of treatment IC_50_ values of CX5461 for TGCT lines ranged from 0.1-1.6 µM (except for GCT72 cells, 3.8 µM), indicating marked potency of CX-5461 against TGCT cells. This is consistent with data in other cancers (ovarian cancer: 12 nM-5.17 µM (15); breast cancer: ∼1.5-11.35 µM (13); colorectal cancer: 0.285-0.835 µM (23, 24); neuroblastoma: 0.2 µM (36); cervical cancer: ∼0.87-0.97 µM (12)). Of note, the control fibroblasts used here displayed an IC50 > 16 µM (Figure 1, Supplementary Figure 3, Table 1). This suggests that (i) the concentrations applied to TGCT lines are in a range comparable to other cancer entities and (ii) nonmalignant cells are markedly less susceptible to CX-5461.

We observed that CX-5461 induces robust, dose- and time-dependent G4 stabilization in TGCT cells, with the strongest effects in EC lines followed by CC and SE cell lines, indicating that TGCT cells (particularly EC) (Figure 2 a-f) are highly susceptible to CX-5461. These data are comparable to results in BRCA1/2 deficient tumors (17), advanced solid tumors (53), glioma (74). By contrast, CX-5461 did not measurably alter BG4 signal in fibroblasts in the tested conditions, underscoring a pronounced G4-targeting effect in TGCT cells under the conditions tested.

In addition, CX-5461 caused a dose dependent decrease in 47S pre rRNA in TGCT cell lines, most strongly in NT2/D1, suggesting inhibition of RNA polymerase I dependent rRNA synthesis by this dual Pol I inhibitor and G4 stabilizer. Similar effects were reported in ovarian cancer (15), uterine leiomyosarcoma (75), multiple myeloma (76), leukemia and lymphoma (44). In contrast, 47S pre-rRNA expression was relatively less reduced in MPAF cells, indicating lower sensitivity of nonmalignant cells to CX-5461 mediated Pol I inhibition and supporting preferential Pol I/G4 targeting in TGCT cells.

Furthermore, we found that CX-5461 induces pronounced, dose- and time-dependent double strand DNA damage in TGCT cells, with the strongest effects in EC lines. Similar damage has been reported in breast (13, 17, 21), neuroblastoma (33), glioblastoma (41), ovarian (14, 15, 32) cancers. In contrast, fibroblasts showed no detectable γH2AX induction under the identical conditions, indicating that CX-5461 induced DNA damage is largely restricted to TGCT cells under the conditions tested.

Cell-cycle profiling revealed that CX-5461 treatment leads to a significant, dose-dependent accumulation of all TGCT cell lines in the G2/M phase, consistent with reports in leukemia (42), ovarian (14, 15, 31), breast (13, 17) cancers. In contrast, fibroblasts showed no detectable change in cell cycle distribution under the same conditions, further supporting a pronounced G2/M arrest in TGCT cells. Moreover, CX-5461 triggered a marked, dose- and time-dependent induction of apoptosis in TGCT cells (again strongest in EC lines), consistent with findings in breast (13, 17, 20), leukemia (42, 53), ovarian (14, 15, 31), neuroblastoma (32, 35), glioblastoma (40). Whereas fibroblasts remained largely unaffected under the same conditions, suggesting that CX-5461 preferentially induces apoptosis in TGCT cell.

Consistently, in line with the finding in leukemia and lymphoma (43, 44, 77), breast (13, 21), ovarian (15), osteosarcoma (28) cancers, transcriptomic profiling revealed a robust activation of p53 signaling pathways following CX-5461 treatment in TGCT cells, including induction of canonical p53 target genes such as CDKN1A, MDM2, PIDD1, GADD45A, ZMAT3, SESN1 and PPM1D, which are involved in cell-cycle arrest, DNA damage responses and apoptosis. Notably, unlike many other malignancies, such as ovarian, uterine carcinosarcoma, lung, breast and colorectal cancers, where TP53 is mutated in >80% of cases (78), somatic TP53 mutations are extremely rare in TGCTs (79). We conclude that CX-5461 exerts its cytotoxic effects in TGCTs predominantly via activation of an intact p53 pathway. Notably, p53 signaling pathway genes harbor predicted G4 regions in their promoters and gene bodies, which should be experimentally explored to address the contribution of G4s in modulating p53 pathway specific responses in TGCT cells.

CX-5461 is a well characterized G4 stabilizing ligand in clinical trials for advanced hematologic malignancies and solid tumors, where it has shown promising antitumor activity (10, 17, 31, 53, 54). It stabilizes G4 structures, impairs replication fork progression and inhibits rRNA synthesis by disrupting RNA polymerase I recruitment, thereby inducing nucleolar stress and activating p53 dependent cell fate programs such as cell cycle arrest and apoptosis. In TGCT cells, we observed DNA damage, G2/M arrest and apoptosis, consistent with these p53 linked CX-5461 responses.

In summary, this study demonstrates that CX 5461 stabilizes G4 structures, disrupts RNA polymerase I transcription and induces DNA damage, leading to activation of p53 dependent pathways, G2/M arrest and apoptosis in TGCT cells. Its minimal impact on non-malignant cells supports its potential as a therapeutic candidate for TGCTs. These findings lay the groundwork for future translational studies and support the continued development of CX-5461 as a treatment option for TGCTs, particularly in patients with chemo-resistant or refractory disease. Ongoing and future efforts should investigate whether CX-5461 can restore cisplatin sensitivity. Collectively, this work advances our understanding of G4 biology in TGCTs and positions CX-5461 as a promising candidate for clinical translation in the TGCTs treatment.

## Supporting information

Supplementary excel sheets

## Acknowledgements

We thank Greta Zech for excellent technical assistance. We also thank the Flow Cytometry Core Facility (Institute of Molecular Medicine and Experimental Immunology, University Hospital Bonn) and the NGS Core Facility (Medical Faculty, University of Bonn/West German Genome Center, WGGC) for their support and access to instrumentation.

## Author Contributions

HS and RI designed the study with expert input from KP. RI, HS and KP contributed to the conceptual development of the project. CR assisted with the optimization of BG4 flow cytometry experiments, and AL assisted with the optimization of western blot analyses. RI performed all experiments. RI and HS drafted the manuscript, and RI, HS, and KP conducted the final critical revision. All authors read and approved the final manuscript.

