## Supplementary Table for "G-quadruplex targeting by CX-5461 as a novel therapeutic strategy in testicular germ cell tumors"

**Supplementary Table S4: IC<sub>50</sub> values (μM) for TGCT and control cell lines following BRACO19 treatment.** Results are presented for 24, 48, and 72 h post-treatment. Color coding denotes potency: red (0-16 μM, very high potency), blue (>16-35 μM, high potency), Yellow (>35-72 μM, moderate potency), and white (>72 μM, low potency).

| Entity | Cell line | IC50 (μM) |  |  |
| --- | --- | --- | --- | --- |
|  |  | 24 h | 48 h | 72 h |
| Embryonal carcinoma | NT2/D1 | 20.17 | 19.32 | 6.065 |
|  | 2102EP | 55.81 | 22.64 | 11.12 |
|  | NCCIT | 19.20 | 63.63 | 14.61 |
| Seminoma | TCam2 | 79.27 | 30.00 | 15.98 |
| Choriocarcinoma | JAR | 68.93 | 70.64 | 8.373 |
|  | JEG3 | 58.13 | 34.60 | 12 |
| Yolk sac tumor | GCT72 | 37.89 | 31.86 | 30.46 |
| Fibroblast | MPAF | 94.60 | 23.90 | 15.77 |

**Supplementary Table S5: IC<sub>50</sub> values (μM) for TGCT and control cell lines treated with CM03 at 24, 48 and 72 h post-treatment.** Results are color-coded by potency: red (0-0.2 μM, very high potency), blue (>0.2-0.5 μM, high potency), yellow (>0.5-1 μM, moderate potency) and teal (>1 μM or unstable, low potency).

| Entity | Cell line | IC50 (μM) |  |  |
| --- | --- | --- | --- | --- |
|  |  | 24h | 48h | 72h |
| Embryonal carcinoma | NT2/D1 | 0.5392 | 0.2606 | 0.1220 |
|  | 2102EP | 0.2519 | 0.1761 | 0.1625 |
|  | NCCIT | 0.2695 | 0.1236 | 0.05255 |
| Seminoma | TCam2 | 0.8039 | 0.2114 | 0.0001362 |
| Choriocarcinoma | JAR | 4.353 | 0.1264 | 0.1242 |
|  | JEG3 | 1.389 | 0.2898 | 0.1877 |
| Yolk sac tumor | GCT72 | 0.8795 | 0.8212 | 0.4936 |
| Fibroblast | MPAF | Unstable | Unstable | 1.106 |

**Supplementary Table S6: IC50 values (μM) for TGCT and control cell lines following PDS treatment.** Results are shown for 24, 48, and 72 h post-treatment. Color coding denotes potency: red (0-10 μM, very high potency), blue (>10-20 μM, high potency), Yellow (>20-30 μM, moderate potency), and Teal (>30 μM, low potency).

| Entity | Cell line | IC50 (μM) |  |  |
| --- | --- | --- | --- | --- |
|  |  | 24h | 48h | 72h |
| Embryonal carcinoma | NT2/D1 | 9.949 | 9.550 | 6.360 |
|  | 2102EP | 20.18 | 18.71 | 10.4 |
|  | NCCIT | 17.44 | 8.497 | 7.881 |
| Seminoma | TCam2 | 24.03 | 20.12 | 10.36 |
| Choriocarcinoma | JAR | 32.69 | 3.940 | 3.053 |
|  | JEG3 | 37.27 | 17.68 | 3.252 |
| Yolk sac tumor | GCT72 | 18.29 | 13.75 | 10.88 |
| Fibroblast | MPAF | 195.8 | 15.27 | 10.89 |

**Supplementary Table S7: IC50 values (μM) for TGCT and control cell lines following RHPS4 treatment, determined using the XTT viability assay.** Results are presented for 24, 48, and 72 hours post-treatment. Color coding denotes potency: red (0-3 μM, very high potency), blue (> 3-10 μM, high potency), Yellow (>10-20 μM, moderate potency), and Teal (>20 μM-Unstable, low potency).

| Entity | Cell line | IC50 (μM) |  |  |
| --- | --- | --- | --- | --- |
|  |  | 24h | 48h | 72h |
| Embryonal carcinoma | NT2/D1 | 2.727 | 0.7771 | 0.1733 |
|  | 2102EP | 28.60 | 1.990 | 1.853 |
|  | NCCIT | 8.636 | 1.997 | 1.671 |
| Seminoma | TCam2 | 7.864 | 4.686 | 3.865 |
| Choriocarcinoma | JAR | 5.760 | 2.583 | 2.053 |
|  | JEG3 | 5.838 | 2.223 | 1.923 |
| Yolk sac tumor | GCT72 | 11.10 | 12.51 | Unstable |
| Fibroblast | MPAF | 78.25 | 51.19 | 5.955 |

**Supplementary Table S8: IC50 values (μM) for TGCT and control cell lines following CX-3543 treatment, determined using the XTT viability assay.** Results are presented for 24, 48, and 72 hours post-treatment. Color coding denotes potency: red (0-4 μM, very high potency), blue (>4-10 μM, high potency), Yellow (>10-20 μM, moderate potency), and Teal (>20 μM, low potency).

| Entity | Cell line | IC50 (μM) |  |  |
| --- | --- | --- | --- | --- |
|  |  | 24h | 48h | 72h |
| Embryonal carcinoma | NT2/D1 | 3.723 | 2.045 | 1.804 |
|  | 2102EP | 5.311 | 3.381 | 5.470 |
|  | NCCIT | 6.528 | 1.398 | 3.656 |
| Seminoma | TCam2 | 1.239 | 1.165 | 1.007 |
| Choriocarcinoma | JAR | 6.469 | 6.354 | 3.517 |
|  | JEG3 | 10.93 | 32.48 | 40.56 |
| Yolk sac tumor | GCT72 | 3.230 | 3.3291 | 3.584 |
| Fibroblast | MPAF | 3.738 | 2.093 | 31.26 |

**Supplementary figure 1: Double-strand DNA break analysis using γH2AX.** **a.** Flow cytometric detection of double-strand DNA damage: gating on single cells (FSC-A vs FSC-H), followed by separation of live and dead cells using Zombie NIR viability dye vs SSC-A; γH2AX levels were quantified as MFI in live cells. **b.** Confirmation of γH2AX staining specificity, with background levels defined by unstained and γH2AX-ve controls. **c.** Western blot analysis of γH2AX further confirmed CX5461-induced DNA damage in NT2/D1 cells treated with CX5461 (1 μM or 2 μM) for 24 h and 48 h. β-Actin served as a loading control.

**Supplementary figure 2: BG4 signal quantification using flow cytometry.** **a.** Flow cytometric quantification of BG4 signal: gating on single cells (FSC-A vs FSC-H), followed by separation of live and dead cells using Zombie NIR viability dye vs SSC-A; BG4 levels were quantified as MFI in live cells. **b.** Confirmation of BG4 staining specificity, with background levels defined by unstained and BG4-ve controls. **c.** Confirmation of increased BG4 signal in NCCIT cells treated with CX-5461 (1 μM, 48 h).

**Supplementary figure 3: Nonlinear regression analysis of CX-5461 sensitivity and $IC_{50}$ calculation.** Four-parameter nonlinear regression curves were used to calculate $IC_{50}$ values for CX-5461 treatment at 24 h, 48 h and 72 h. Curves are derived from XTT cell viability data presented in Figure 1.

**Supplementary figure 4: Nonlinear regression analysis of BRACO19 dose-response curves and $IC_{50}$ determination.** BRACO19 reduces the viability of TGCT (**a-g**) and fibroblast cells (**h**). XTT cell viability assays ($n = 5$) were performed in TGCT cell lines and non-malignant control fibroblasts treated with 2, 5, 10, 20, 30 and 50 μM BRACO19 for 24, 48 and 72 h. BRACO19 exposure resulted in a significant, concentration- and time-dependent decrease in TGCT cell viability, with a similar cytotoxicity profile observed in MPAF cells. DMSO vehicle controls were included at each concentration and set to 100% viability; data are normalized to the corresponding vehicle controls and presented as mean ± SD. $IC_{50}$ values for BRACO19 treatment at 24, 48, and 72 hours were determined by fitting four-parameter nonlinear regression curves to XTT cell viability data.

**Supplementary figure 5: Nonlinear regression analysis of CM03 dose-response curves and IC<sub>50</sub> determination. a-g.** CM03 reduces viability of TGCT cells. XTT cell viability assays (n = 5) were performed in TGCT cell lines and non-malignant control fibroblasts treated with 0.05, 0.10, 0.25, 0.50 and 1 μM CM03 for 24, 48 and 72 h. CM03 exposure resulted in a significant, concentration- and time-dependent decrease in all TGCT cell viability, whereas MPAF cells (**h**) were not or only minimally affected. DMSO vehicle controls were included at each concentration and set to 100% viability; data are normalized to the corresponding vehicle controls and presented as mean ± SD. IC<sub>50</sub> values for CM03 treatment at 24, 48 and 72 h were determined by fitting four-parameter nonlinear regression curves to XTT cell viability data.

**Supplementary figure 6: Nonlinear regression analysis of PDS dose-response curves and IC<sub>50</sub> determination.** PDS reduces viability of TGCT (**a-g**) and control fibroblast cells (**h**). XTT cell viability assays (n = 5) were performed in TGCT cell lines and non-malignant control fibroblasts treated with 2, 5, 10, 15, 20 and 25 μM PDS for 24, 48 and 72h. PDS exposure resulted in a significant, concentration- and time-dependent decrease in viability across all TGCT cell lines, with a similar level of cytotoxicity observed in MPAF cells (**h**). DMSO vehicle controls were included at each concentration and set to 100% viability; data are normalized to the corresponding vehicle controls and presented as mean ± SD. IC<sub>50</sub> values for PDS treatment at 24, 48 and 72 h were determined by fitting four-parameter nonlinear regression curves to XTT cell viability data.

**Supplementary figure 7: Nonlinear regression analysis of PRHPS4 dose-response curves and IC<sub>50</sub> determination.** RHPS4 reduces viability of TGCT (**a-g**) and control fibroblast cells (**h**). XTT cell viability assays (n = 5) were performed in TGCT cell lines and non-malignant control fibroblasts treated with 0.5, 1, 2, 3.5, 5, and 10 μM RHPS4 for 24, 48 and 72h. RHPS4 exposure resulted in a significant, concentration- and time-dependent decrease in viability across all TGCT cell lines, whereas MPAF cells (**h**) were not or only minimally affected up to 48 h but showed a similar cytotoxic effect at 72h. DMSO vehicle controls were included at each concentration and set to 100% viability; data are normalized to the corresponding vehicle controls and presented as mean ± SD. IC<sub>50</sub> values for RHPS4 treatment at 24, 48 and 72 h were determined by fitting four-parameter nonlinear regression curves to XTT cell viability data.

**Supplementary figure 8: Nonlinear regression analysis of CX-3543 dose-response curves and IC<sub>50</sub> determination.** CX-3543 reduces viability of TGCT (a-g) and control fibroblast cells (h). XTT cell viability assays (n = 5) were performed in TGCT cell lines and non-malignant control fibroblasts treated with 0.5, 1, 2, 3, 5, and 10 μM CX-3543 for 24, 48 and 72h. CX-3543 exposure resulted in a significant, concentration- and time-dependent decrease in viability across all TGCT cell lines, as well as MPAF. DMSO vehicle controls were included at each concentration and set to 100% viability; data are normalized to the corresponding vehicle controls and presented as mean ± SD. IC<sub>50</sub> values for CX-3543 treatment at 24, 48 and 72 h were determined by fitting four-parameter nonlinear regression curves to XTT cell viability data.

Supplementary figure-1

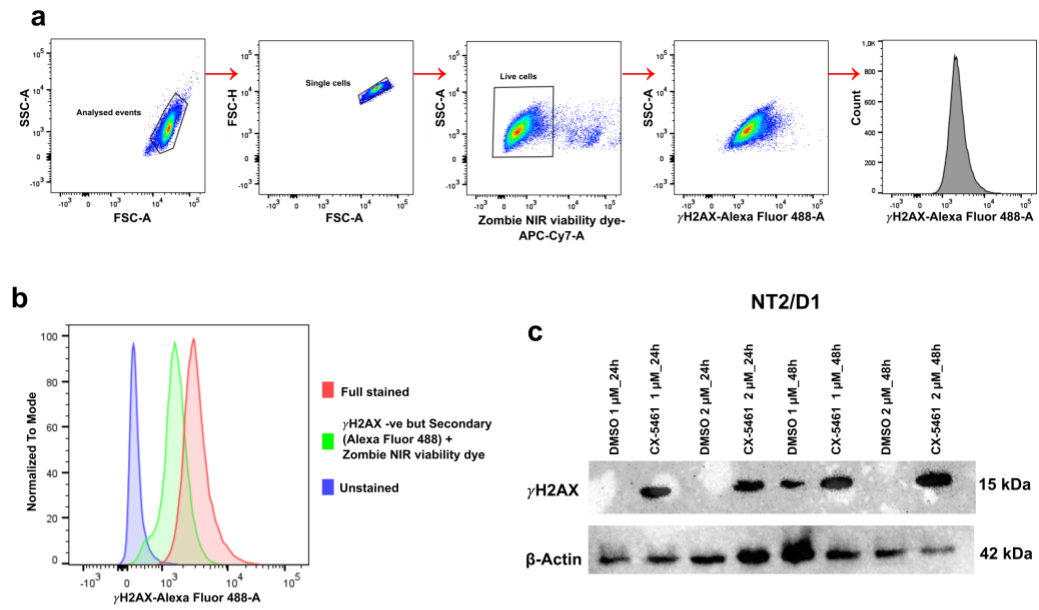

Supplementary figure-2

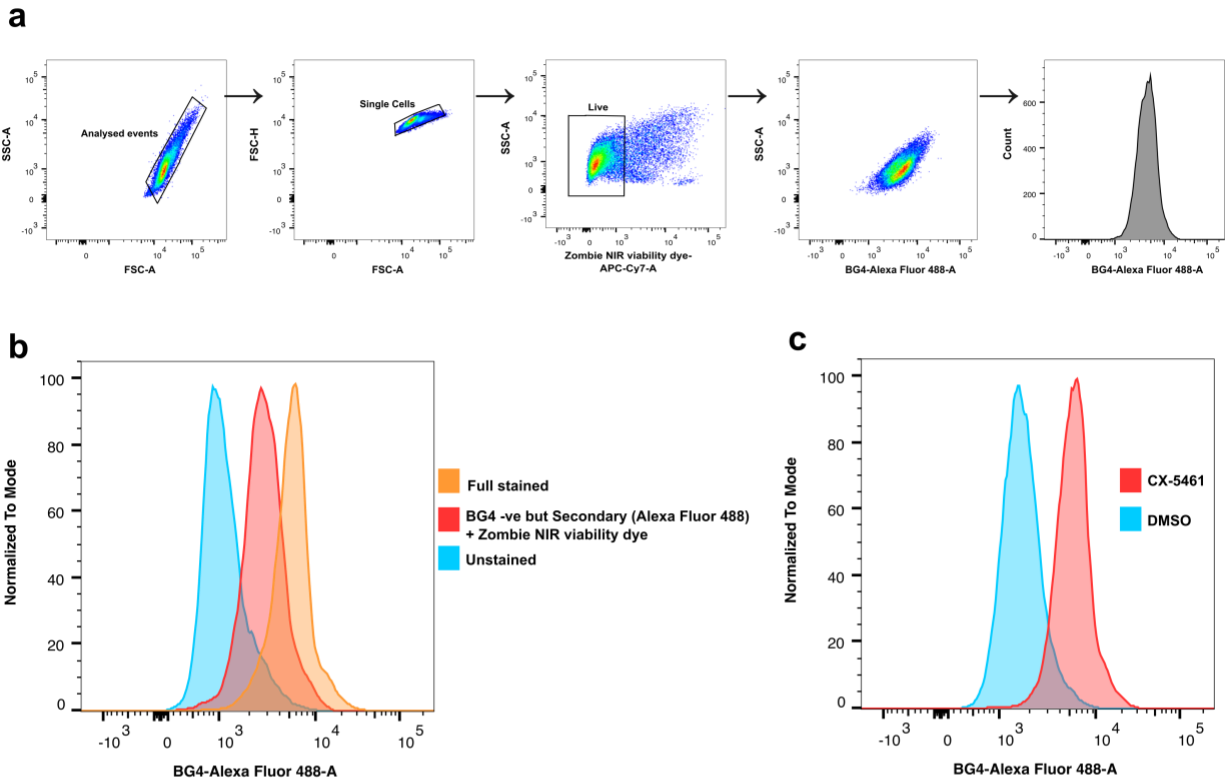

Supplementary figure-3

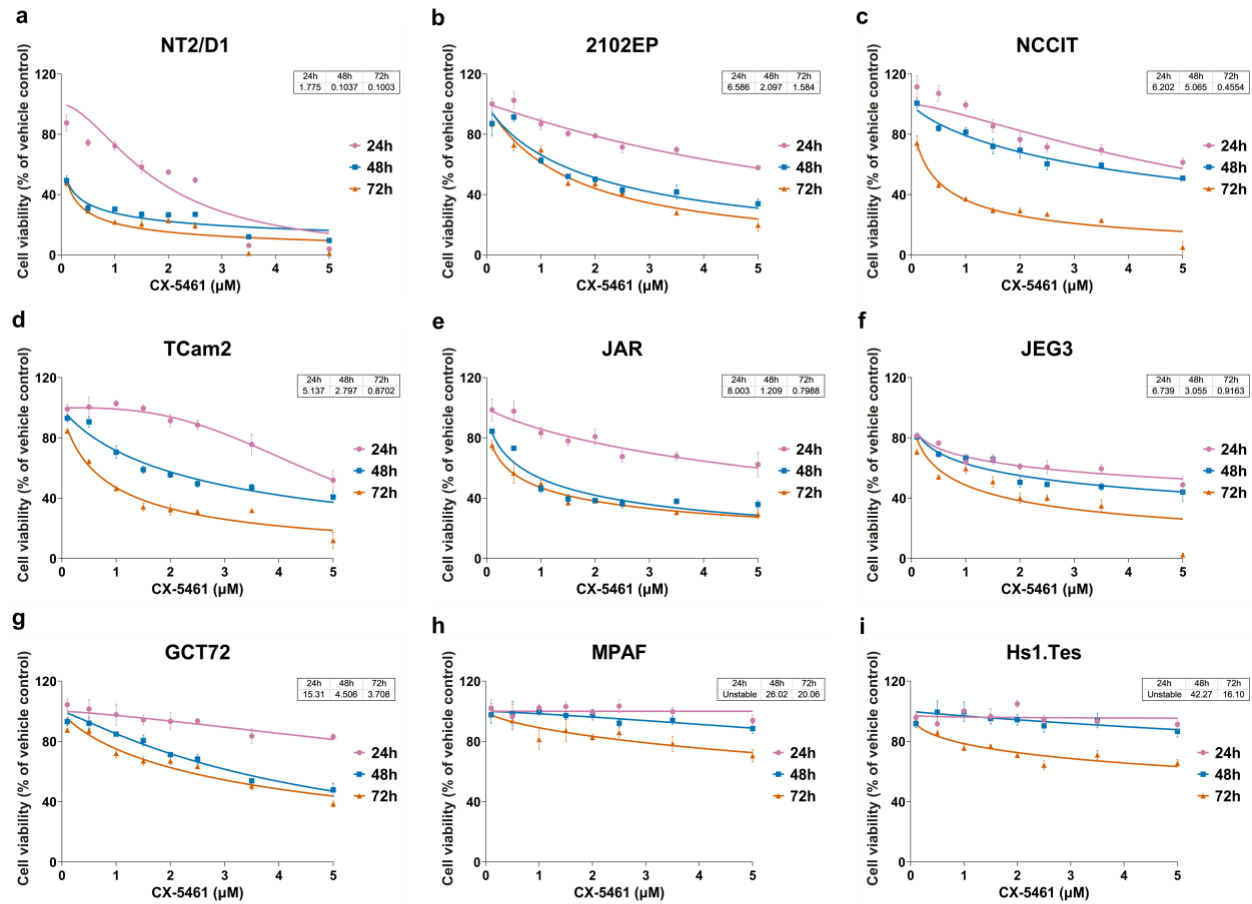

Supplementary figure-4

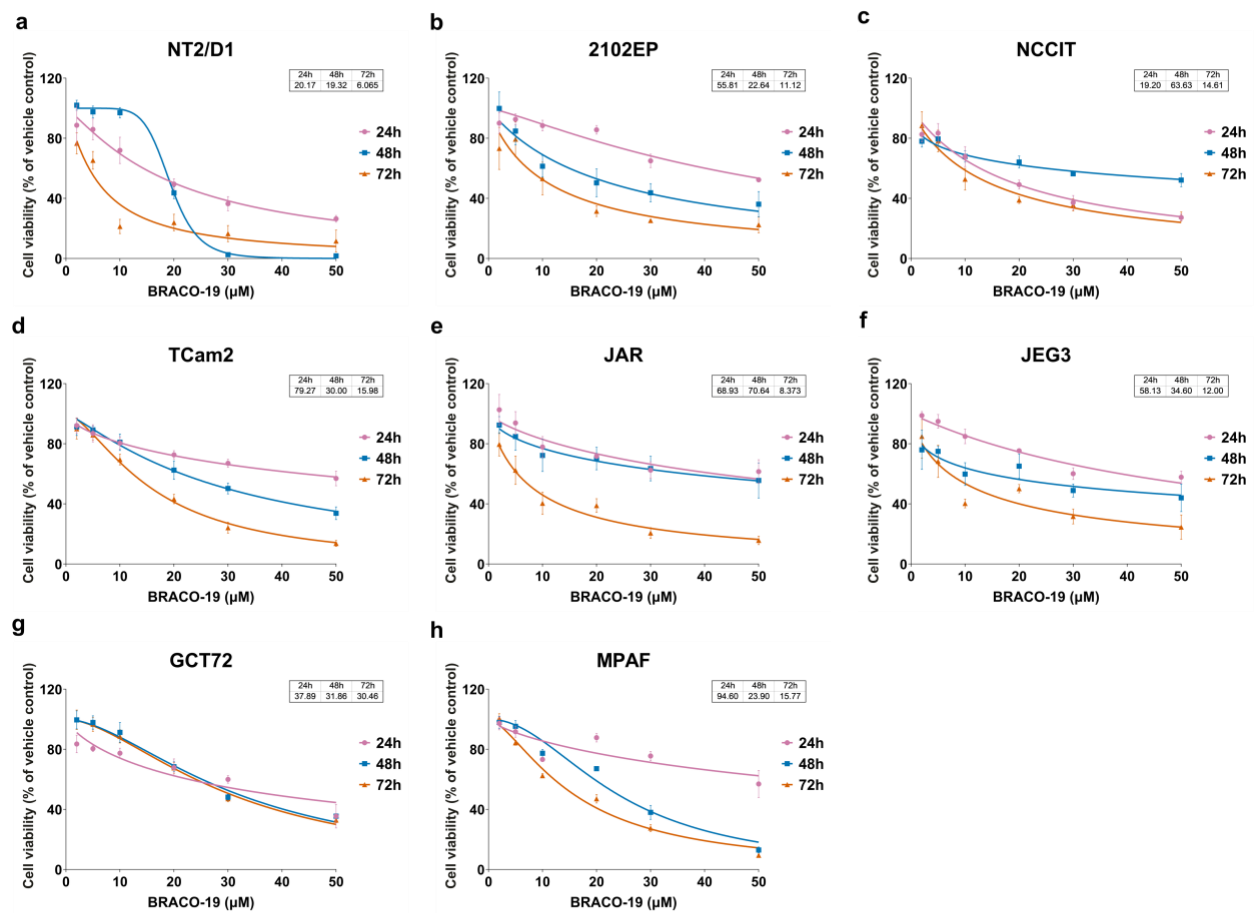

Supplementary figure-5

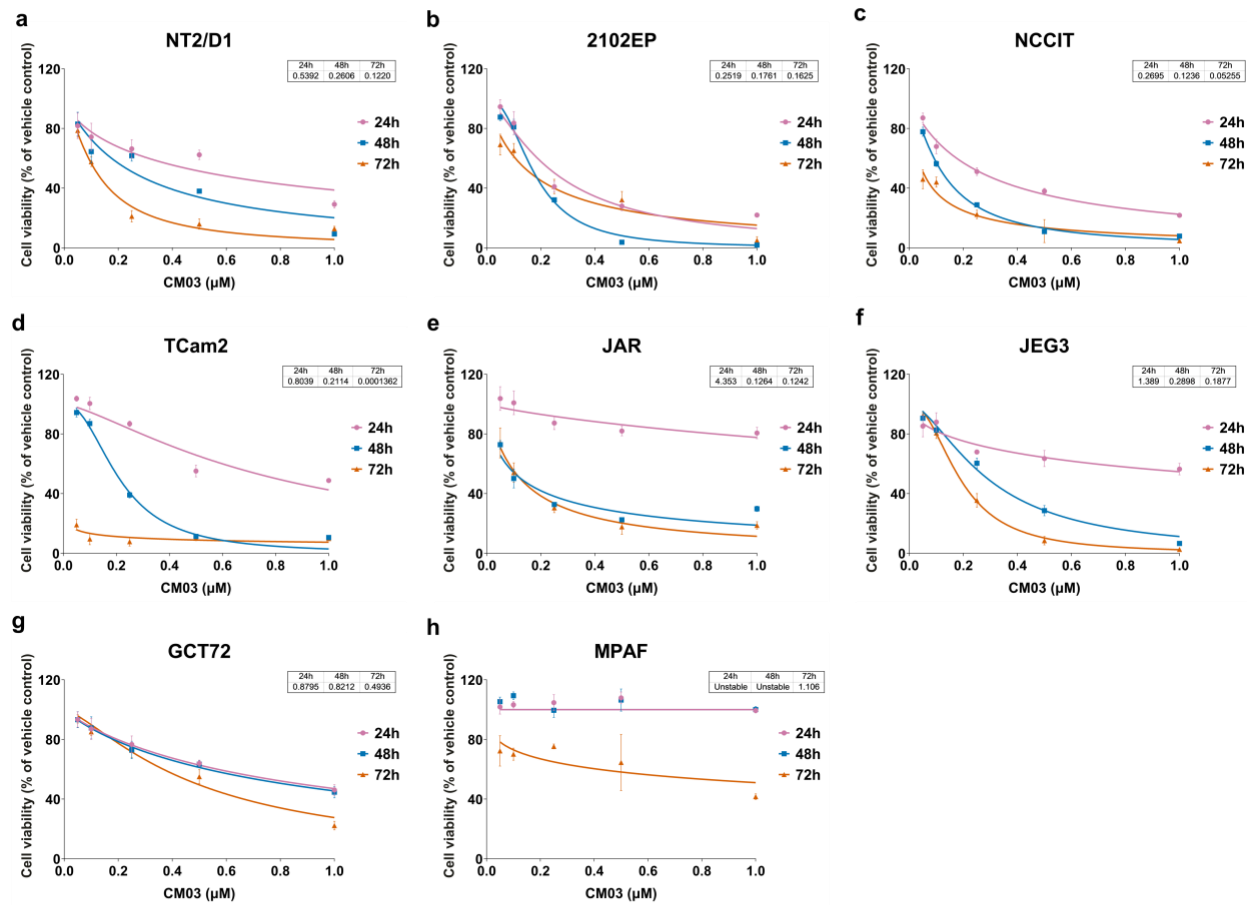

Supplementary figure-6

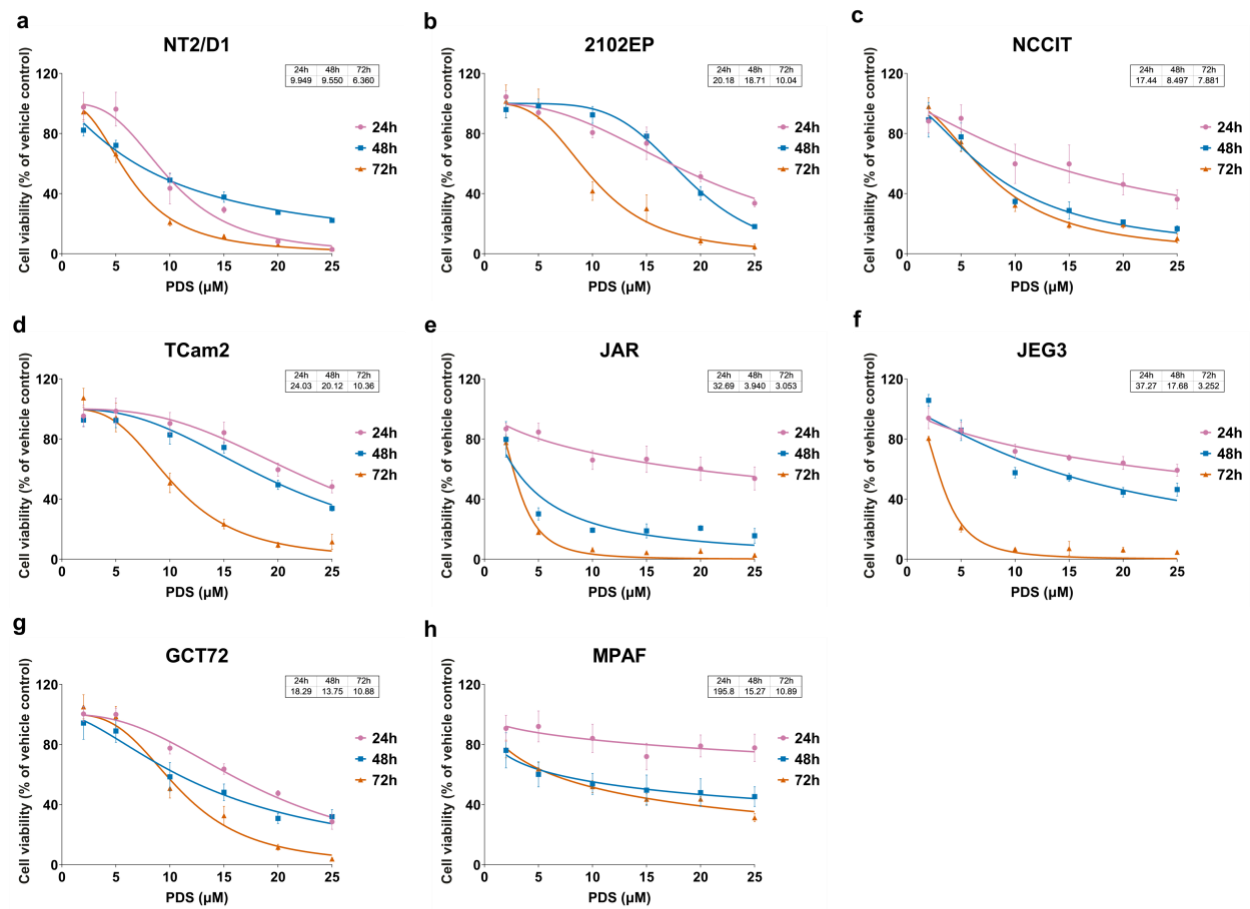

Supplementary figure-7

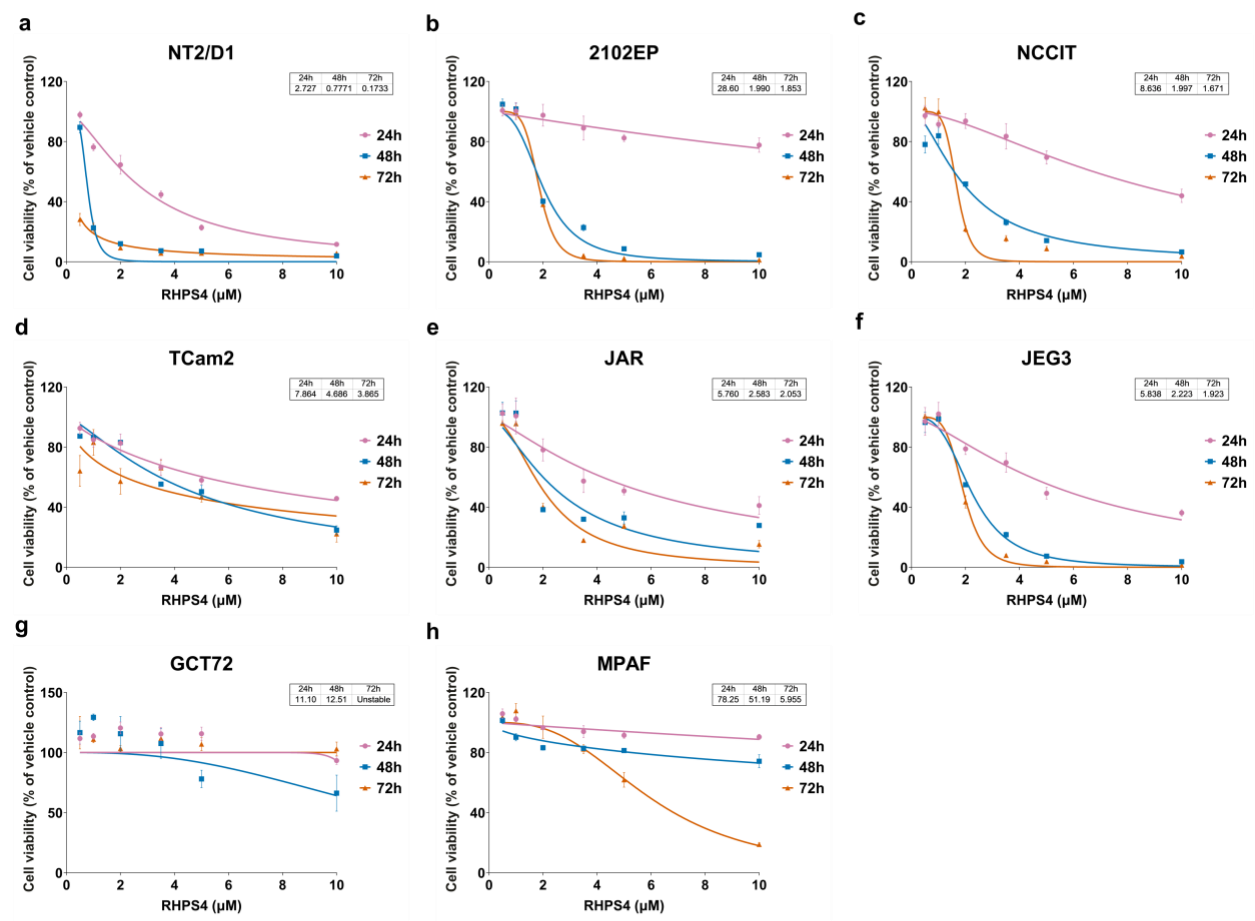

Supplementary figure-8

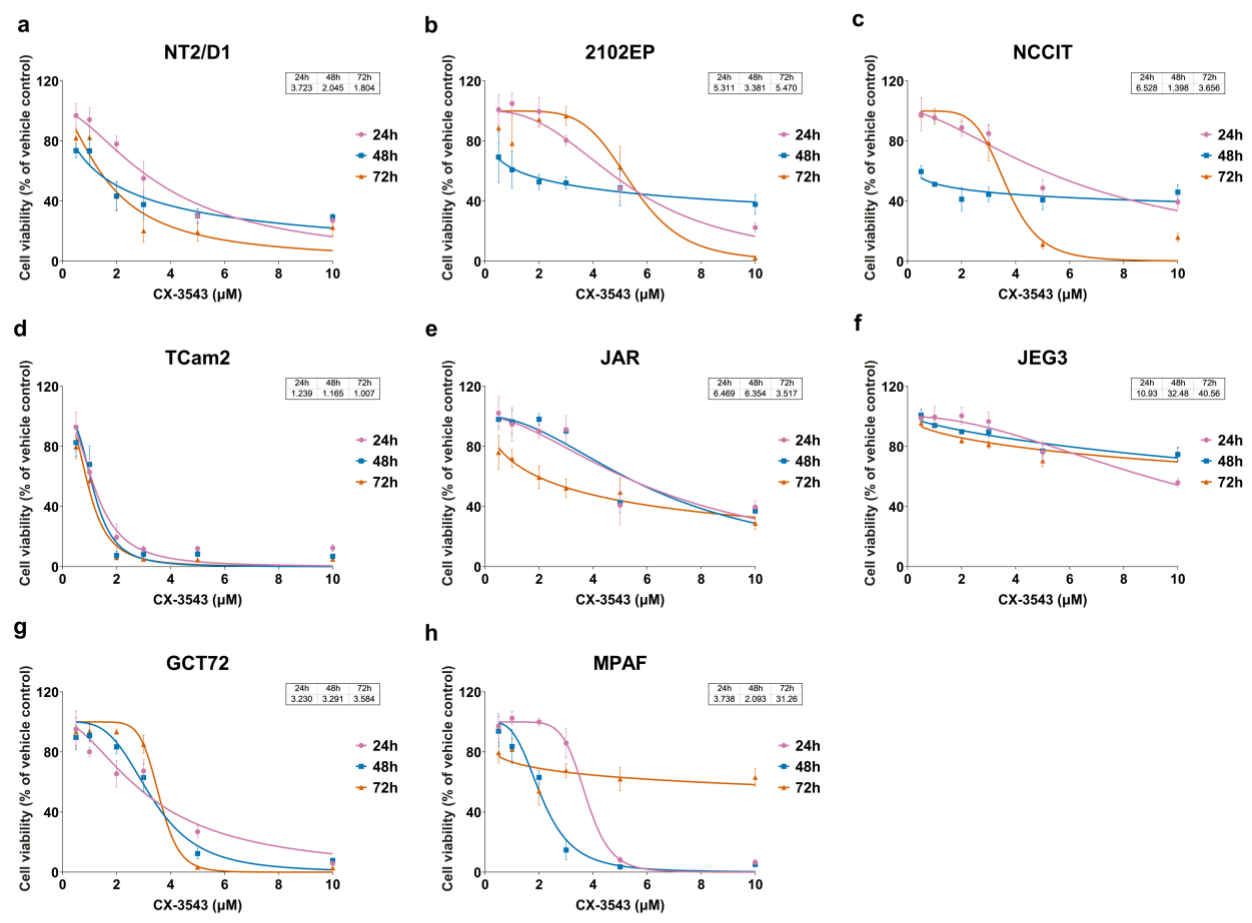
